# Sand throwing in a pit-building antlion larva from a soil mechanical perspective

**DOI:** 10.1101/2020.11.25.398073

**Authors:** Sebastian Büsse, Thies H. Büscher, Lars Heepe, Stanislav N. Gorb, Hans Henning Stutz

## Abstract

Sandy pitfall traps are an elaborate construction to capture prey and antlions are well-known representatives of this predation technique. From a soil mechanical perspective, antlions exploit the interactions between the particles of their habitat and engineer a stable trap. This construction is close to the unstable state, where a prey item will immediately slide towards the center - towards the ambushing antlion - when accidently entering the trap. This method is efficient, but requires permanent pit maintaining. According to the present knowledge, antlions throw sand at their prey, to distract it, and/or cause sand slides towards the center of the pit. Using sand throwing and escape experiments, as well as finite element analysis, we supported this hypothesis. Furthermore, we added new hypothesis about maintaining the pitfall trap. We showed that sand that accumulates in the center of the pit will be continuously removed, which lead to the slope maintenance close to an unstable condition. This avoids self-burial of the antlion, as well as decreasing the chance of prey item escapes by keeping the slope angle steep. This demonstrates the interaction of an insect larva with its abiotic environment from a novel perspective and adds further insights into longstanding entomological hypotheses.

## Background

Trap-building is a highly specialised, but comparably uncommon, hunting strategy within the animal kingdom (Franks et al. 2019). Most trap-building invertebrates employ silk in their constructions, with orb-web spiders probably being the most prominent example (Denny 1976; Vollrath and Knight 2001). These invertebrates successfully use silk to capture prey (Lin et al. 1995; Krink and Vollrath 2000; Venner et al. 2006). In contrast, the construction of traps without the employment of silk is best known in wormlions (Diptera: Vermileonidae) and antlions (Neuroptera: Myrmeleontidae), using sand to dig a pitfall trap (Fig. 1 B,C) (cf. Adar et al. 2016) with a few exceptions (cf. Dejean et al. 2005). Even though the trap-digging strategy in antlions (spiral digging) is considered more effective in comparison to central digging in wormlions (Tuculescu 1975; Franks et al. 2019), pitfall trap building strategies of worm- and antlions represent an excellent example for convergent evolution of behaviour (Miler et al. 2018).

**Figure 1:**
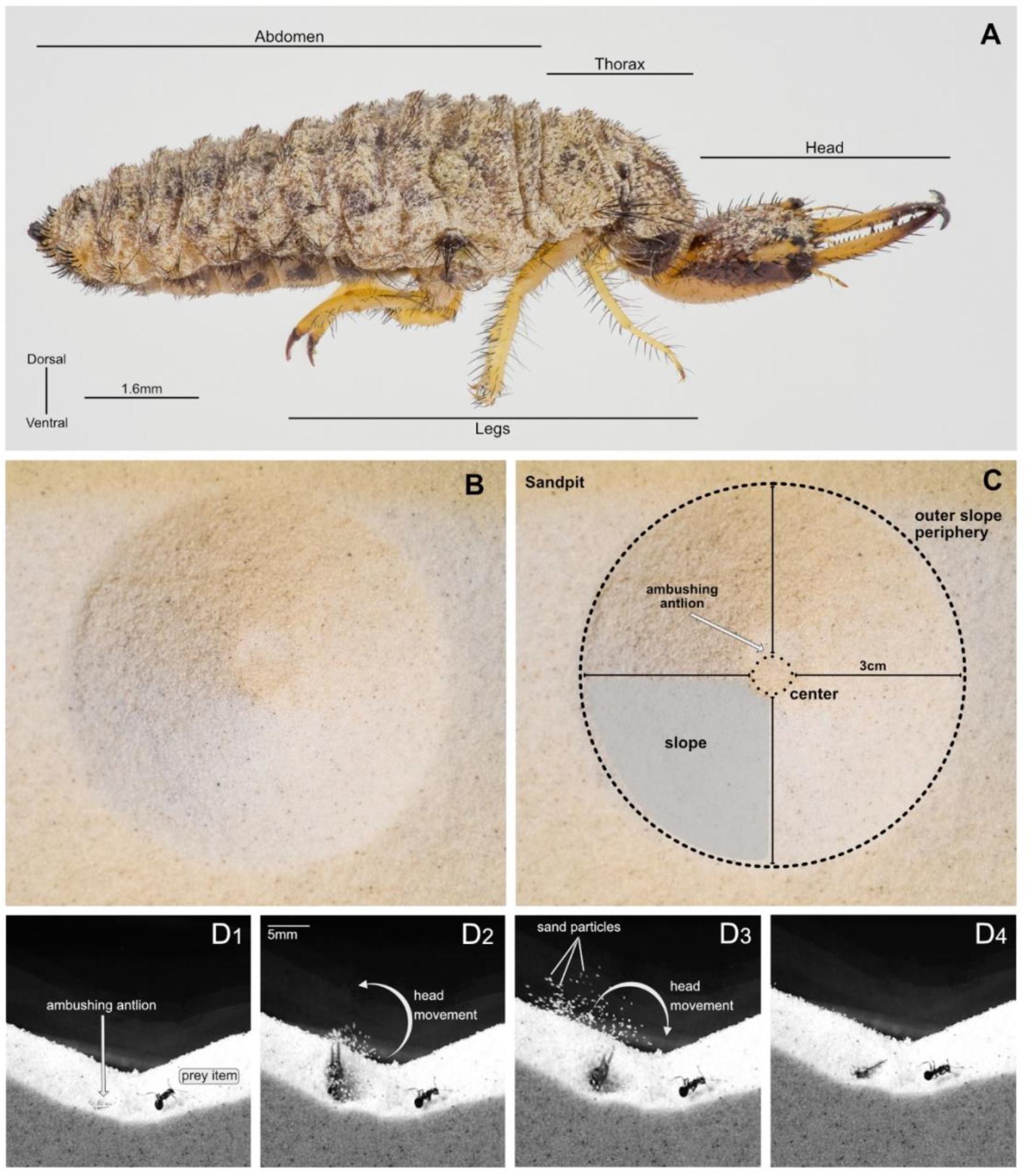
Antlion and pitfall trap. A. Habitus of antlion larva (*Euroleon nostras*), lateral view. B-C. Sandpit used from the antlion as pitfall trap. C. Same sandpit as in B including labels. D. Sand throwing behaviour of the antlion by the flick of its head. D1-4. Time series based on single frames obtained from high-speed videography: 1. Before the flick of the antlion head starts; 2. During the flick, upwards movement of the antlions head; 3. During the flick, downward movement of the antlions head; 4. After the flick of the antlion head.

Generally, ground-dwelling animals which inhabit sandy habitats are strongly affected by the physical characteristics of the substrate. Sand is a collection of particles interacting with each other via contact forces. Here, spontaneous organisation (Rosato et al. 1987; Möbius et al. 2001) faces natural stratification (Bak et al. 1987; 1988), which leads to an irregular solid-pore system (Herrmann 1998). Whereas the number of pores in granular media influences the stress-deformation behaviour to a great extent, loose packing of granular particles leads to an instable construct that can easily reach an unstable state (Terzaghi 1943, Miura et al. 1997). Here more voluminous sand grains show a larger angle of repose (definition see “Methods: Terminology”) than less voluminous grains, and all interact with the successive sand layers (Makse et al. 1997).

The juvenile stages of most antlion species (Myrmeleontidae) utilise exactly this soil mechanics phenomenon. The laval antlion (Fig. 1 A) is a pit-building ambush predator (Franks et al. 2019). The hunting success of it largely depends on the abiotic factors of its habitat (Scharf and Ovadia 2006; Bar-Ziv et al. 2019), such as sand grain size and distribution, and the majority of antlion species prefer sand with a comparably small particle size (Allen and Croft 1985; Loiterton and Magrath 1996; Botz et al. 2003; Farji-Brener 2003). Suitable substrates will enable the construction of considerably larger pits (Barkae et al. 2012), resulting in the ability to capture larger prey and reducing the risk of prey escapes (Griffiths 1980; Lucas 1982; Heinrich and Heinrich 1984; Scharf et al. 2018). The relationship between sand properties and slope is the key difficulty for prey items captured in a sandy pitfall trap, as shown for the ant species *Aphaenogaster subterranea* (Latreille, 1798). This species copes with the unstable substrate with a gait pattern transition from the tripod gait to the metachronal wave pattern (Humeau et al. 2019). Changing the gait to the one involving a higher number of legs (e.g. metachronal wave) is known from other insects adapting to challenging attachment conditions (e.g. walking on the ceiling), and thus risking to lose their grip to the substrate (Gorb and Heepe 2017; Büscher and Gorb 2019). Thus, the pitfall trap of an antlion is a mechanically unstable construction, and its capturing success is increasing with increasing slope angles and positively affected by decreasing sand particle size (Botz et al. 2003). Exploiting the instability of the slope, the trap’s morphology (Fig. 1 B,C) is used to facilitate hunting prey of very different kind and size (cf. Gepp and Hölzel 1989). The larval antlion is ambushing in the vertex of the pit (Lambert et al. 2011), where it is throwing sand, using fast flicks of its head (Fig. 1 D_1_-D_4_; Griffiths 1980; Gepp and Hölzel 1989; Lambert et al. 2011). This behaviour intends the distraction of escaping prey and more importantly, causes small sand sandslides to trap the prey and translocate it to the center of the vertex - respectively towards the antlion (Griffiths 1980).

We here present a study of the soil mechanical behavior that ensure the antlion’s prey capturing success. Underlining the sandslide theory, mentioned above, on the one hand, we present a supplementary hypothesis focusing on pit maintaining, on the other. Presumably, the sand throwing will not only actively prevent the prey from escaping, but will also maintain the required instability of the slope. The sand, which will accumulate at the center of the pit by the movements of the escaping prey, will be removed and more importantly the slope will be kept close to an unstable condition. This study exemplifies the benefits of an interdisciplinary approach to evaluate a known phenomenon from the perspective of two different scientific disciplines aiding in the understanding of the underlining mechanisms – here of the sand throwing by antlion larvae.

## Methods

### Terminology

#### Angle of repose

Physically, the angle of repose is described as the angle at which a transition between phases of granular materials happening. The adopted common definition is the steepest slope angle of the unconfined granular material measure from the horizontal axis.

#### Friction angle

The friction angle defines the frictional shear resistant of the soil dependent of the normal effective stress.

#### Cohesion

The cohesion is the shear strength component of the soil that is independent of the inter-particle friction.

#### Mobilised friction

This is the definition of the friction that was mobilised in the strength reduction method using the finite element model.

### Sand throwing experiments

Larvae of *Euroleon nostras* (Fourcroy, 1785) were kept in small ant-terrariums (210×100×105mm) filled with sand (particle size: 125 μm). Prior the experiments, the antlion had 24h for setting up their pitfall trap. We used small instars of the house cricket (*Acheta domesticus* (Linnaeus,1758)) as well as black garden ants (*Lasius niger* (Linnaeus,1758) and *L. fuliginosus* (Latreille, 1798)) to film the prey capturing process of the antlion larvae using an Olympus OMD 10mkII digital camera (Olympus K.K., Tokyo, Japan) equipped with a Leica 45mm macro lens (Leica, Wetzlar, Germany). For measuring the slope angles (N=9, total sequences 16) and for further image processing, Affinity Photo and Affinity Designer (Serif Ltd, Nottingham, United Kingdom) were used. The slopes before and after sand throwing were compared via a paired t-test, as the data was normally distributed (according to Shapiro-Wilk’s test for normality, P=0.08), using SigmaPlot 12.0 (Systat Software Inc., San José, CA, USA).

### Escape experiments

For the escape experiments a small formicarium (210×100×105mm) was used to film house crickets (*A. domesticus*) while trying to escape a conical half-shaped artificial pitfall trap. The formicarium was filled with sand (particle size: 125μm) using a defined funnel to produce a pit close to the unstable state. Furthermore, we used ants (*L. niger* and *L. fuliginosus*) to escape from a conical artificial pitfall trap. To produce this pit a box was filled with sand (see above) with a hole in the bottom to produce a pit close to the unstable state. The prey items were filmed, using an Olympus OMD 10mkII digital camera (Olympus K.K., Tokyo, Japan) equipped with a Leica 45mm macro lens (Leica, Wetzlar, Germany), while ascending the slope of the pit (N=7). Furthermore, a house cricked was filmed, using a Go-Pro Hero 5 (GoPro Inc., San Mateo, US) in time lapse setting (1 frame/min), over the course of 12h by trying to escape the pit (respectively the terrarium), without the maintaining of an antlion. For measuring the slope angles and for further image processing, Affinity Photo and Affinity Designer (Serif Ltd, Nottingham, United Kingdom) were used.

### Photography

For stacked photography, we used a custom-made 3D-printed LED illumination dome system (Bäumler et al. 2020) and an Olympus OMD 10mkII digital camera (Olympus K.K., Tokyo, Japan), equipped with a Leica 45mm macro lens (Leica Camera AG, Wetzlar, Germany). In general, all images were subsequently processed in Affinity Photo and Affinity Designer (Serif Ltd, Nottingham, United Kingdom).

### Finite-element simulations

The simulations are conducted using the finite-element method (FEM; commercial software package OPTUM G2 2020, Copenhagen, Denmark). For studying the slope stability, the strength reduction method has been applied to simulate the progressive failure of the sand slope that is built by the antlion larvae as pitfall trap. The underlying principle of the strength reduction method is that the initially assigned soil strength parameters will be reduced until a failure occurs in the soil continuum. A detailed description of the strength reduction method is given in e.g. Tschuchnigg et al. (2015a; 2015b). In short, the strength reduction method is used to estimate the stability of a soil mechanical system by reducing systematically the strength parameters of soil, namely cohesion and the friction angle. For the assessment of the failure, the factor of safety (FoS) is used as:

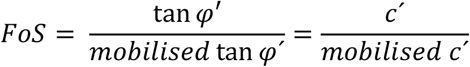

Where *φ′* is the effective friction angle of the soil and *c′* is the effective cohesion of the soil. Herein, the friction angle and the cohesion describe the shear strength of a soil using the concept of the Mohr-Coulomb failure criterion (Mohr 1900). The friction angle defines the friction shear resistant of the soil dependent of the normal effective stress. The cohesion is the shear strength component of the soil that is independent of the interparticle friction. These are divided by mobilised friction and cohesion. The mobilised friction tan\phi and cohesion describe the values that could be applied in the strength reduction method. FoS < 1 describe a failure and FoS > 1 describe a stable pit slope. The axisymmetric geometry of the initial reference pit is shown in Figure 2. The boundary conditions at the bottom of the model are fixed for all degree of freedoms, and the right side is a slider boundary condition.

**Figure 2:**
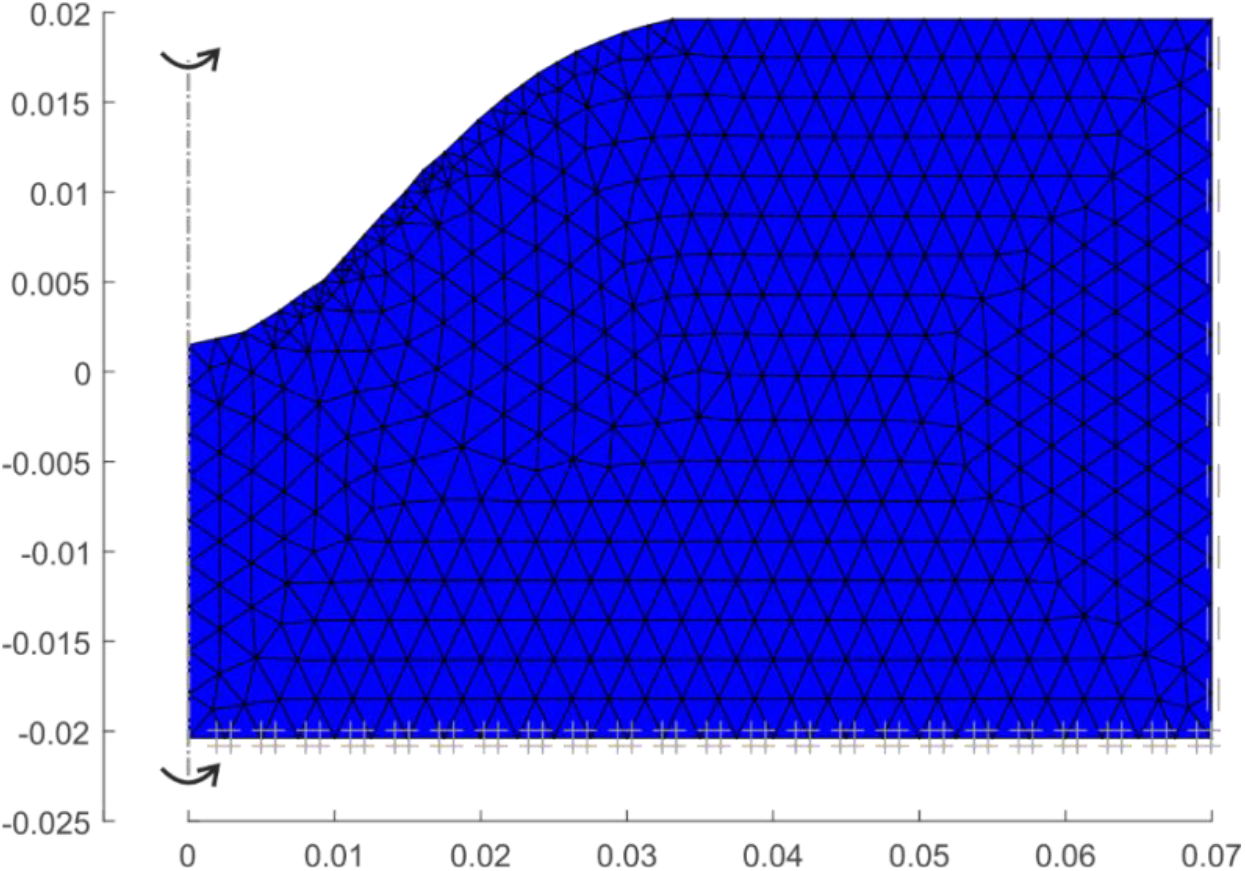
Axi-symmetric mesh for the initial slope model, using 1000 (15-noded) elements. The soil (blue) is modelled using the linear elastic – perfectly plastic Mohr-Coulomb model.

The model used in the simulations is the linear elastic-perfectly plastic Mohr-Coulomb constitutive model, which have been proven to be sufficient for such ultimate limit state simulations (Davis 1968, Tchuchnigg et al. 2015). The used geometry is chosen based on the previously reported average antlion pits (e.g. Bongers & Koch 1981; Lucas 1982) and our experimental setup.

In general, the following steps are conducted for each simulation:

1. The initial stress is applied and calculated.
2. The initial geometry is analysed to estimate the initial FoS.
3. The changed geometry is used to estimate the change in the FoS and the consequences due to the sand throwing of the antlion larvae.

The reference configuration (Fig. 2) has an initial FoS= 1.096, and this means the slope geometry is stable. For the more accurate prediction of the failure mechanisms and the FoS, a mesh adaptivity step is applied with three adaptive iterations using the shear dissipation as adaptivity control. The mesh adaptivity is a procedure using an adaptive meshing technique to refine the mesh around the shear zone in which the plastic deformation is overdriven (Ortiz & Quigley 1991). The mesh is refined according to the norm of the strain vector ||ε||. The initial model shown in figure 2, shows a mesh consisting of 1000 elements; here, the model is using a linear elastic – perfectly plasticity Mohr-Coulomb constitutive model (Mohr 1900). The parameters used in the simulation are given in Table 1. In the mesh refinement step, 2000 elements are used.

**Table 1:** Parameters used in the finite-element simulations for the Mohr-Coulomb model (grey zones do not influence the strength reduction method)

| Material | Cohesion<br>c [kPa] | Friction<br>angle $\varphi$ [°] | Dilatancy<br>[°] | E-modulus<br>[MPa] | Poisson's ratio<br>[-] |
| --- | --- | --- | --- | --- | --- |
| Sand initial | 0 | 34.5 | 0 | 20 | 0.2 |
| Sand reduced density | 0 | 29 | 0 | 20 | 0.2 |

In total, six different simulations were conducted. In these simulations, the slope geometry was changed to simulate the throwing behaviour of the antlion larvae (Case 1 & 2). In two simulations, the soil was simulated with reduced friction angle zones (Case 3 & 4), based on the looser soil state. This looser state is based on the assumption of generation of looser soil zones due to the sand throwing behavior. This was done to screen the effect, when there is no volume loss in the sand. In the last two simulations, the change in geometry (sand throwing) and change in density was applied (Case 5 & 6). Based on the sand throwing experiments and the experimental observations. The modelling assumption here is that a looser granular packing has a smaller angle of friction (Mitchell & Soga 2005). The changes in the geometry and the changes in the areas with smaller friction angles are indicated in Table 2.

**Table 2:** Change in model areas to simulate the six different case via strength reduction method

| Case | Area of slope geometry<br>change [mm<sup>2</sup>] | Area of change in<br>friction angle [mm<sup>2</sup>] |
| --- | --- | --- |
| 1 | 4.64 | - |
| 2 | 3.40 | - |
| 3 | - | 15.66 |
| 4 | - | 8.19 |
| 5 | 4.64 | 12.13 |
| 6 | 3.40 | 11.48 |

This selection aims to model the different effects induced on the soil by the antlions sand throwing behaviour, to study the effect of the antlion trap/pit slope stability. The figures were prepared with MATLAB (R2019b, The Mathworks Inc., Natick Massachusetts) using the data files form OptumG2.

All experiments and simulations were conducted considering a dry sand character, the effect of partially saturation of the soil was not studied.

## Results

We used the insights from biological experiments considering the soil mechanical properties of the antlions’ trap building and combined these with finite element simulations to identify the underlying soil mechanical behaviour.

### Sand throwing experiments

After allowing the antlion larvae to set up a pitfall trap for 24h, all formicariums for the experiment exhibited a sandpit ready to capture prey. After inserting a prey item into the terrarium, the antlions start throwing sand (Fig. 1, supplemental videos 1-3), when noticing the vibrations of the prey. The sand throwing can start without visible sand movement, but becomes more frequent (sand throwing and therefore sand movement), when the prey item changes the slope geometry and especially when moving sand from the slope towards the center of the pit (respectively towards the ambushing antlion). The sand throwing of the antlion usually causes small sand slides (supplemental video 1) distracting the prey and/or causing the prey sliding towards the center of the pit (supplemental video 2). However, it becomes obvious that these sand slides also recover the steepness of the sandpit’s slope - smoothened by the movement of the prey or the antlion itself (Fig. 3, supplemental video 3). The average slope angle before sand throwing of the antlion is 27.3 +/- 2.7 ° (min. 22.5°, max. 31.5°), the average slope angle after sand throwing of the antlion is 31.1 +/- 2.1° (min. 26.5°, max. 34.5°) resulting in an average slope angle change of 3.44° (min. −1°, max. 7°). The sand throwing of the antlion results in a significantly steeper slope after the action (paired t-test, t=-8.095, d.f.=8, N_1,2_=9, P≤0.001).

**Figure 3:**
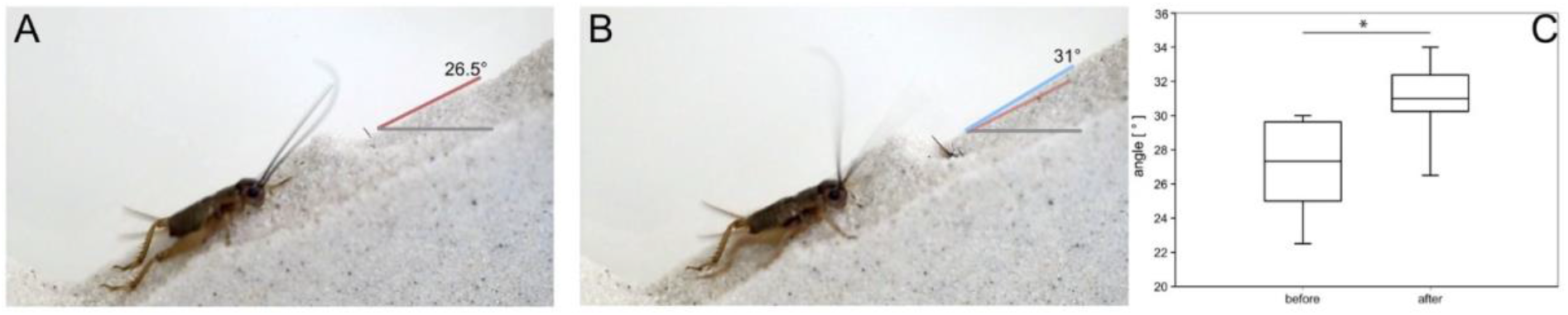
Slope of a sandpit before and after sand throwing by the antlion. A. Already leveled slope (caused by the prey item, house cricket on the left side) before the sand throwing of the antlion. Red line indicates the slope angle of 25.5°, the grey line indicates the reference angle. B. Slope after the sand throwing of the antlion. Blue line indicates the new slope angle of 31°, red line of the old slope angle in transparent, the grey line indicates the reference angle. C. Box-whisker-plots of initial (before) and resulting (after) slope angles of the pit. The line represents the median, the box and whiskers the 10, 25, 75 and 90 % percentiles, respectively. * P≤0.001, paired t-test.

### Escape experiments

The restructuring of the sand topography within the artificial sandpits reveals the influence of the distortion, a prey item causes without the influence of maintaining the pit by the antlion (Fig. 4, supplemental video 4 and 5). The escape efforts of the prey item can cause serious damage to the sandpit geometry, especially to the slope angle. The slope in the 12h experiment decreases from 38° over 29° after 6h, to 21° after 12h (Fig. 4, supplemental video 4). However, even single events (one walk of a cricket or ant on the slope) can cause sand movements and therefore changes in the overall slope geometry. Sand is pushed downwards, towards the canter of the slope by every step of the prey item. Every step is, therefore, changing the slope geometry slightly, as well as pushes small volumes of sand to the centre of the pit. The influence of several steps of the prey item accumulates over time. Consequently, with an increasing dwelling time of the prey in the pit, the change of the sandpit geometry leads to an increasing chance of its escape (supplemental video 5).

**Figure 4:**
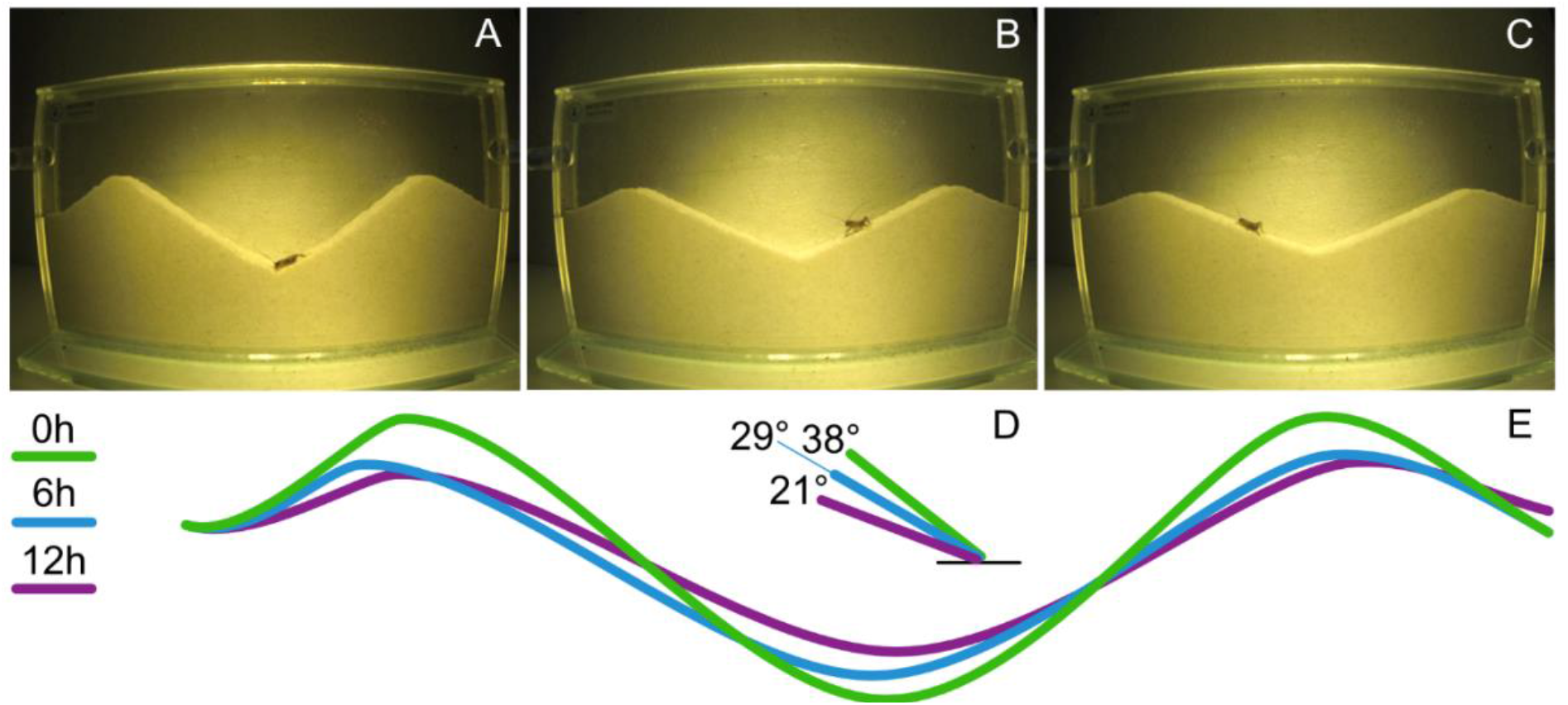
Escape experiment: house cricket over the cause of 12h in a formicarium. A-C. Change of the artificial pit geometry over the cause of 12h. A 0h. B 6h. C 12h. D. Change of the slope angle over the cause of 12h, green line after 0h, blue line after 6h and violet line after 12h. E. Change of the artificial sandpit geometry over the cause of 12h shown as schematic, green after 0h, blue after 6h and purple after 12h.

### Finite-element analysis

The results of the conducted finite element analysis are shown in Figure 5. The first two initial cases (case 1 and 2; Fig. 5 A,B, see also Table 2) demonstrate the effect of a factor of safety (FoS) reduction, based on a slight change in the slope geometry. Depending on this geometry change (case 2, Figure 5 B), the FoS can result in a value below 1.0, which indicate an unstable slope (initial slope geometry FoS = 1.096). This change in the slope angle could be observed in the sand throwing experiments (see Figure 3 A, B). Herein, a local change of the slope geometry can generate an instable slope. In figure 5 (C,D), the results of the cases 3 and 4 with a pure change in density are simulated without a change in the slope geometry. Here it can be seen that a change in density may be caused by the movement and throwing behavior of the ambushing antlion and can lead to an unstable condition.

**Figure 5:**
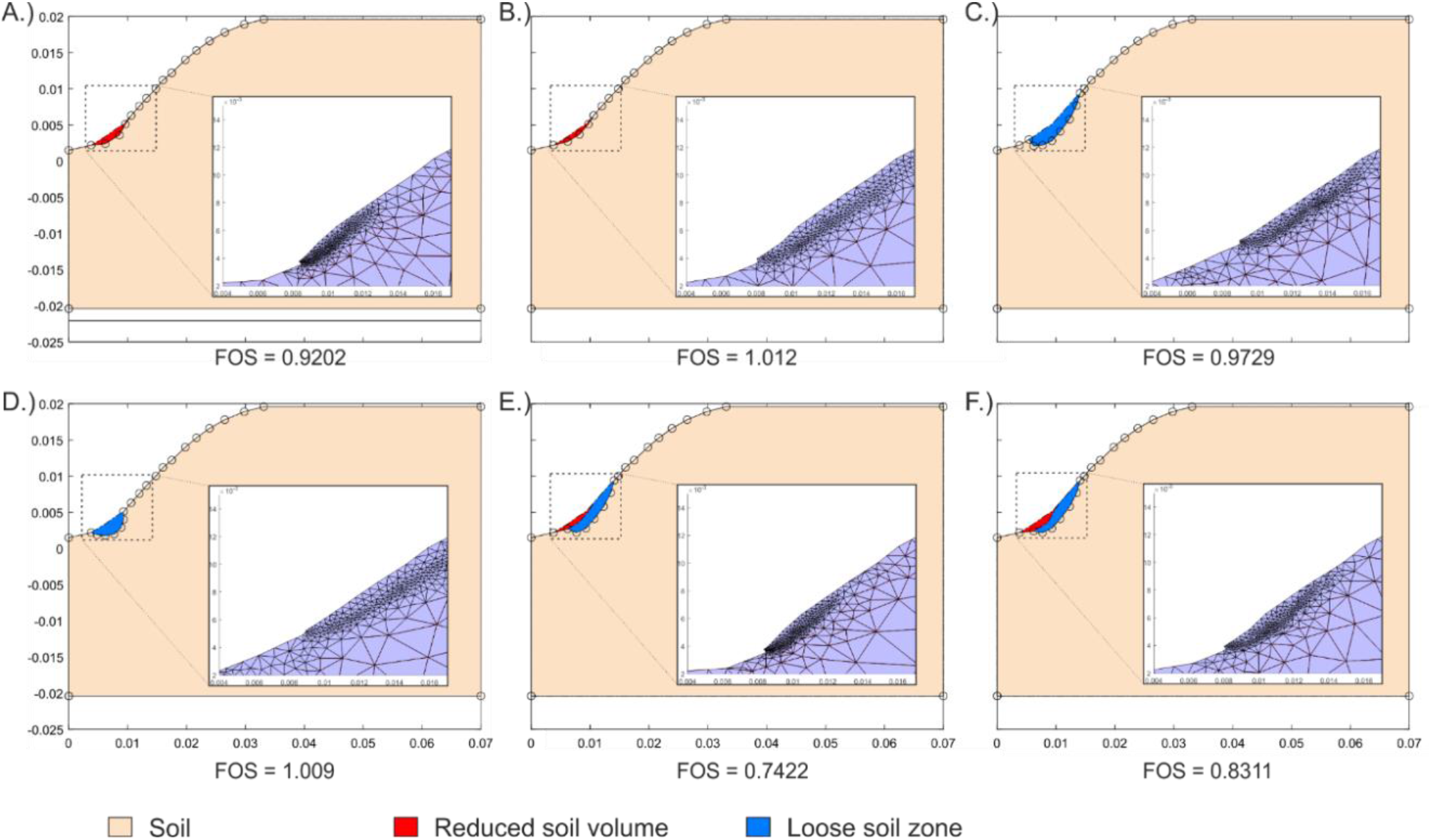
Results for the Cases 1 – 6 (A.-F.) with the indicated change of the reduced soil volume (in red), the changed zones for the friction angle (in blue) and the results, shown as failure surfaces with different adaptive meshes, which demonstrate the failure mechanisms.

Generally, local changes do not necessarily lead to an unstable situation (see Fig. 5 D; FoS, 1.009). Whereas, a combination (cases 5 and 6) of a change in the slope geometry (case 1 and 2; Fig. 5 A,B) with a change in density of the soil (represented by a change in the friction angle; case 3 and 4) leads to an unstable condition. Cases 5 and 6 (Fig. 5 E,F) are the most realistic natural scenarios compared to the described sand throwing experiments. In both cases, the FoS is below 1.0, which indicates an unstable slope.

Besides an adaptive remeshing was used to refine the mesh around plastic zones in the soil. The use of the remeshing technique lead to small element sizes close to zones of localized deformations (shear zones). Therefore, the meshes shown in figure 5 indicate the different failure geometries and shear zones, which are similar to the geometries in the sand throwing experiments (Figure 3 and supplementary video material).

## Discussion

The results of the sand throwing and escape experiments are combined with the finite element simulations to underline the resulting hypothesis as well as add a new soil-mechanical hypothesis. The sand throwing behaviour of antlion larvae is used during pit building (Bongers & Koch 1981) as well as prey capturing (Griffiths 1980). During pit building, the antlion sorts the sand grains towards a preferably smaller grain size (Allen and Croft 1985; Loiterton and Magrath 1996; Botz et al. 2003; Farji-Brener 2003) by the sand throwing, which allows for larger and more stable pits (Barkae et al. 2012). Further, during prey capturing, the sand throwing is used to cause small sand slides that displace the prey item towards the ambushing antlion (Griffiths 1980). However, a prey item can cause significant structural damage to the pit’s geometry (cf. escape experiments). Therefore, pit maintaining is vital for the antlions prey capturing success. Since antlions usually built their pits close to the natural equilibrium condition of the slope (Botz et al. 2003) given by the angle of repose of the granular media (Allen and Croft 1985; Loiterton and Magrath 1996; Botz et al. 2003; Farji-Brener 2003), the pit’s slope is highly unstable and delicate to disturbances (Lucas 1982). Here, without the maintenance by the antlion, the slopes are unstable and the prey causes an irreversible deformation to the slope angle (slope angle reduction; Fig. 4). Therefore, without constant maintaining of the pitfall trap (during prey contact), the antlion befalls self-burial and the slope angle shallows (Fig. 4), so that a prey item can more easily escape. However, the capturing success is increasing with an increasing slope angle causing a prey item more likely to slide towards the center of the pit (Botz et al. 2003). As indicated in the sand throwing experiment, the slope inclination increases by the sand throwing behaviour of the antlion leading to retaining an unstable state (Fig. 3), the fact highly supporting the pit maintaining hypothesis.

From a soil mechanical perspective, the soil state is changing from looser to a denser state as the thrown sand causes a reorganization of the particles along the slope of the trap. The sandparticles sliding towards the center of the pit are rearranged during this relocalisation and come to rest in a denser conformation. The finite element simulation supports our observations and experiments, because only a combined mechanism (cases 5 and 6, change in density and sand volume) brings the slope to an unstable state from an initial stable one (mean slope angle change of 3.44°, Fig. 3), as the factor of safety (FoS) of 0.74 (case 5) and 0.83 (case 6) clearly shows. On the other hand, the finite element simulations underline the previous hypothesis that the sand throwing causes small sandslides (Griffiths 1980), as also shown in the supplemental videos (1 and 2). Here the change of FoS under 1 (unstable state) in the simulation is indicating that the sand indeed slides towards the center of the pitfall trap. Additionally, sand slides may provide the information to the sensory system of the predator about an optimal repose angle of the pit.

## Conclusions

We challenged the prevailing hypothesis on antlion sand throwing by investigating the mechanism with a combination of sand throwing observations, escape experiments as well as finite element simulations. Our results support the existing hypothesis that small sand slides displace the prey item towards the ambushing antlion (Griffiths 1980), but furthermore add a soil mechanical perspective to this behaviour: pitfall traps of antlion larvae are mechanically unstable constructions, where the prey capturing success increases with an increasing slope angle. We show that a prey item can considerably change the slope geometry (flatten the slope) in the course of 12 h (if no antlion is involved; see. Fig. 4). Furthermore, the sand throwing experiments reveal significantly higher slope angles after the sand throwing (if a prey item is involved; see Fig. 3). We hypothesize, that sand throwing functions as the trap maintenance mechanism, to keep the critical slope angle and counteracts self-burial of the antlion itself.

## Ethics

Insect specimens used in this study are not protected, and no ethical statement is necessary.

## Data accessibility

All raw data will be uploaded to Dryad

## Authors’ contributions

SB, LH, SNG and HHS designed the project and developed the concept of the study. THB and SB reared the antlions. SB and THB performed the highspeed video recordings and the experiments. SB and THB analysed the biological experiments and performed the statistics. HHS performed the finite element analysis. HHS and LH formulated the physical principles and analysed the finite element analysis. SB, THB, SNG and HHS wrote the manuscript. All authors agree to be held accountable for the content therein and approve the final version of the manuscript.

## Acknowledgements

We are grateful for the support of the members of the Functional Morphology and Biomechanics Group at Kiel University, especially J. Heepe for providing the samples. We could not accomplish this study without the help of Jonathan Neumann (University of Potsdam). The discussions with Prof. Dr. Franz Tschuchnigg are highly appreciated by HHS.

## Supplement

Supplementary Video 1: Sandslides caused by antlion

Supplementary Video 2: Sandslides caused by antlion and relocalisation of prey item

Supplementary Video 3: Slope change caused by prey item with antlion

Supplementary Video 4: Slope change caused by prey item without antlion over the cause of twelve hours (12h experiment)

Supplementary Video 5: Ant escaping a antlions pitfall trap, without active antlion

